# Outcome of Crash Course Training on Protein Structure Prediction with Artificial Intelligence

**DOI:** 10.1101/2022.09.01.506222

**Authors:** D. Balamurugan, Maureen Dougherty, Joseph Lubin, Paul Arias, Janet Chang, Kenneth Dalenberg, Vlad Kholodovych, Ehud Zelzion, Sagar D. Khare, James Barr von Oehsen, Michael E. Zwick, Stephen K. Burley

## Abstract

Protein structure predictions have broad impact on several science disciplines such as biology, bioengineering, and medical science. AlphaFold2[1] and RoseTTAFold[2] are the current state-of-the-art AI methods to predict the structures of proteins with an accuracy comparable to lower-resolution experimental methods. In its 2021 year review, both these methods were recognized as “breakthrough of the year” by Science magazine[3] and “method of the year” by Nature magazine [4]. It is timely and important to provide training and support on these emerging methods. Our crash course “Enabling Protein Structure Prediction with Artificial Intelligence “was conducted in collaboration with domain experts and research computing professionals. The crash course was well received by the community as there were 750 registrants from all over the world. Here we provide the summary of the crash course, describe our findings in organizing the crash course, and explain what preparation steps helped us with the hands-on training.

**CCS CONCEPTS:** Computing methodologies à Machine learning à Machine learning approaches à Bio-inspired approaches

## 1 INTRODUCTION

Over the past 50 years, computational methods based on homology modeling, molecular dynamics simulation, contact maps, and recently deep learning models have been used to predict the structure of the proteins[1, 2, 5–7]. Today, the top prediction methods are AlphaFold2, developed by Google DeepMind, Inc., and RoseTTAFold, developed at the University of Washington/Howard Hughes Medical Institute. Both methods are AI powered. They utilize attention based deep learning networks, which are popular in the natural language models such as BERT and GPT[2]. AlphaFold2 and RoseTTAFold predict the three-dimensional (3D) structure of proteins (atomic coordinates) with accuracies comparable to lower resolution experimental methods without the need of a previously determined 3D experimental structure (template) of an evolutionarily related protein. Due to this advantage, many researchers want to adopt these methods for their ongoing and future research projects.

CASP (Critical Assessment of protein Structure Prediction) is a popular competition in the structural biology community[8]. CASP began in 1994 and is held once every two years. In the most recent CASP held in 2020, formally called CASP14, AlphaFold2 predicted the structure of an unknown protein from the sequence information with remarkable accuracy[9]. RoseTTAFold has shown a similar performance and accuracy with the structure prediction. Science and Nature magazines announced AI powered protein structure predictions made by AlphaFold2 and RoseTTAFold as the breakthrough and the method of the year 2021, respectively[3, 4].

AlphaFold2 and RosettaFold teams published their results around the same time in 2021, and they open sourced their software packages. In addition to open sourcing their code bases, both supported web-services for doing structure predictions. AlphaFold2 team also built a database of precomputed protein structures called AlphaFoldDB[10, 11] that covers almost all the human proteome. The research community working with proteins wanted to have access to these emerging methods and resources. Our research computing professionals working at Rutgers University Office of Advanced Research Computing (OARC) [12] received a handful of queries from the campus researchers on the availability and accessibility of these methods for the campus research computing facility. We heard similar discussions in the Eastern Regional Network (ERN) [13]associated partner universities. Also, there was a discussion thread in the Campus Champion email list from the professionals working at Cleveland Clinic, Cornell, Harvard, and New York University installing and managing the database of AlphaFold2.

Because computational predictions are now able to provide protein structures before the experimental studies begin, computational approaches can help with the early investigations of time critical research questions in molecular biology, medicine, and biotechnology. For example, when the SARS-CoV-2 pandemic started, the structure of the viral spike protein was modeled computationally, based on experimental knowledge of the template homology models based on SARS-CoV-1Spike, before the structure of SARS-CoV-2 spike was available to the community[14, 15]. When the experimental structures, based on cryo-electron microscopy (cryo-EM), were reported, researchers were able to analyze them to understand the impact of amino acid substitutions observed in Variants of Concern. For example, computed structure models of receptor binding domain of Omicron variants were predicted with the computational methods by combining AlphaFold2, RoseTTAFold, and energy models before the experimental structures were released [16].

Researchers working in proteomics, drug discovery, and medicinal chemistry can benefit substantially from a training course on these deep learning methods for protein structure prediction and their applications. However, the challenge in providing such training demands expertise in multiple fields, including structural biology, machine learning methods, molecular modeling, and computational data science. In this article, we describe our experience organizing the collaborative crash course on “Predicting protein structures with AI”. We believe that the challenges, solutions, and outcome of the crash course will benefit the research computing community in organizing other similar collaborative events on emerging research themes.

We acknowledge that conferences such as PEARC, Super Computing, HPC, IEEE, ACS, APS, etc., organize training programs on cutting edge topics. These training events are immensely useful for the participants. However, these events happen once a year and not all the researchers are able to attend these conferences.

Our crash course was conducted in collaboration with the science domain experts and research computing professionals. The Rutgers University Institute of Quantitative Biomedicine (IQB) [17] co-organized the event in collaboration with the OARC[12], the RCSB Protein Data Bank (RCSB PDB) [18], and the ERN [13]. The OARC, the central research computing facility for all the Rutgers campuses, provided the computing resources for the hands-on training. Subject matter experts from Rutgers departments, centers, and external speakers contributed to the crash course.

The crash course attracted ~750 registrants from around the world. Throughout the crash course, participants were actively engaged via the chat and QA features available on the Zoom virtual platform.

Herein, we describe the organization and outcome of the crash course. We start with a brief overview about the scope and organization of the crash course, next discuss the demographics and response from the participants. We summarize the presentations, describe the preparations for the hands-on training, and conclude with thoughts on what makes on-line training events successful.

## 2 SCOPE AND ORGANIZATION OF THE CRASH COURSE

The objective of the crash course was to update the community on the recent developments in using AI methods for predicting protein structures. It is a topic of great interest to both experimental and computational scientists working in fundamental biology, biomedicine, and bioenergy. The crash course was organized to cover the background and as well as the recent applications to get a sense on where the field is headed. It was organized in three thematic sessions. The first session focused on the science and application of machine learning methods for protein structure prediction. The second session showcased examples of applying these new techniques for the current research going on at all Rutgers University campuses. The final session was dedicated to a hands-on tutorial helping beginners learn about these methods and run protein structure predictions using various computational resources, including freely available web and cloud services, and campus or national supercomputing facilities.

The crash course was planned as a virtual event primarily because of the COVID19 pandemic and the Omicron surge. Being a virtual event, it was easy to scale because there were no restrictions on physical resources limiting the number of participants. Later, in the section on preparations for the hands-on training, we point out the challenges associated with the hands-on exercises for supporting many participants from multiple institutions.

### 2.1 Participants Response and Interest in the Event

The event attracted ~750 registrants. Fig. 1 shows the breakdown of registrations during the two months leading up to the event, home institution, profession, gender, and race/ethnicity. As seen in Fig. 1A, there was a steady increase in the number of registrations over the period of eight weeks. Fig 1B shows the breakdown of the registration by profession. About one third of registrants were graduate students, another one third came from the staff and faculty, with the remainder being undergraduates, postdocs, and group leaders. Rutgers topped the list of host institutions (Fig. 1C). This is not surprising since many of OARC users and the IQB affiliated members were deeply interested in the topic. As we mentioned in the introduction section, OARC staff received several queries about the availability of AlphaFold2 and RoseTTAFold. Fig 1D and 1E show the demographics of the registrants. More than 30% of the registrants were females. From the disclosed numbers (approximately 13% of registrants elected not to disclose their race/ethnicity), 13% were of underrepresented ethnic groups.

**Figure 1:**
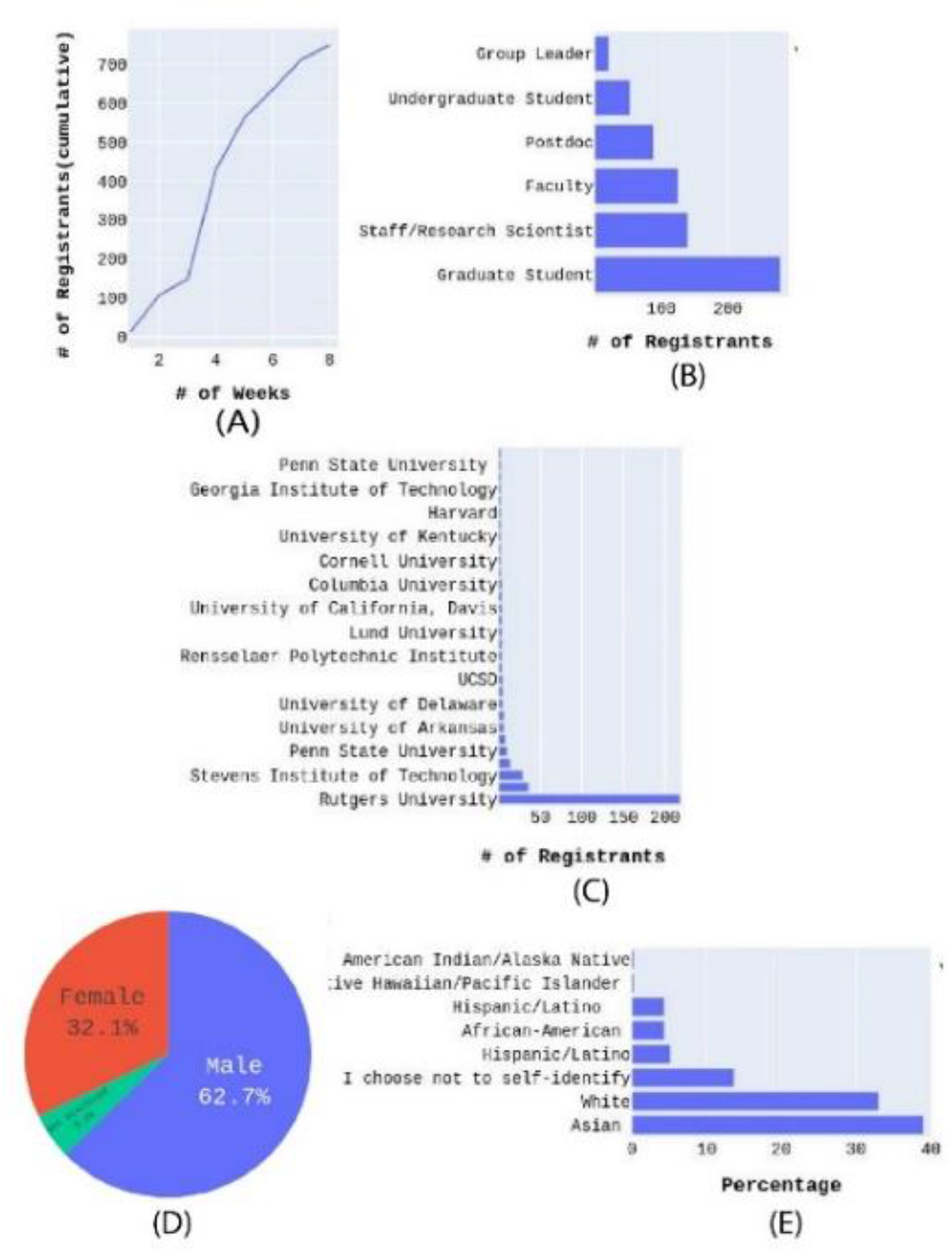
Summary statistics of the registration for the crash course that were collected over eight weeks: (A) the cumulative total number of registrants during the eight-week period, (B) breakdown of registrant by profession, (C) the total number of registrants per university (only top 15 included), and (D) demographics of gender and (E) race/ethnicity

Participants identified themselves as residents of 24 countries (See Fig. 2A). It is worth noting that for some of the Asian countries, the crash course occurred after midnight local time. Throughout the event, participants were actively involved, asking questions in the Q&A feature, posting discussion points in the chat feature in Zoom. We directed participants to use the Q&A feature to post questions for the speakers and post any comments and concerns in the chat feature. We arranged for event support assistance with OARC and IQB staff members, who answered logistic and technical questions in real time. Speakers addressed the questions directed to them or related to their field of expertise.

**Figure 2:**
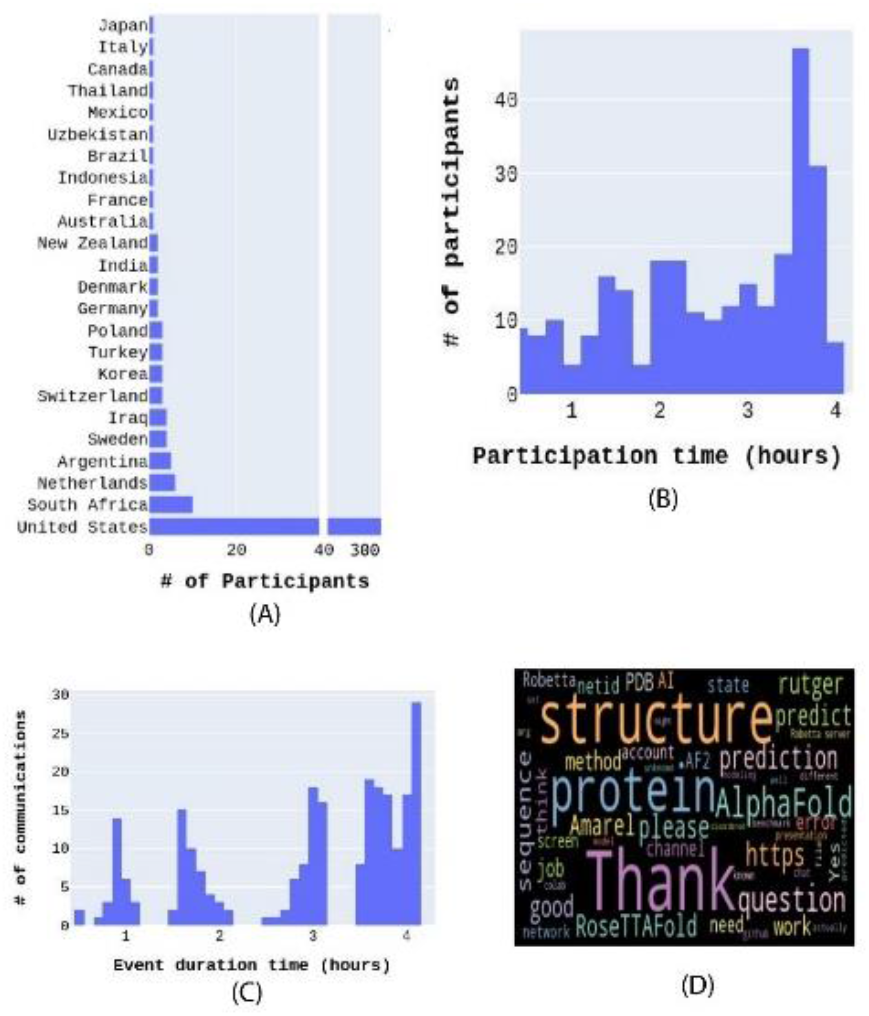
(A) The participants country representation, (B) the histogram of time spent by the participants in Zoom, (C) the number of interactions from all the organizers and participants through chat and Q&A over the course of the event, and the (D) word cloud of the interactions.

Many of the participants attended the entire event. Fig. 2B shows the histogram of the time spent by each participant. We noticed that many had to log in to the Zoom session multiple times. This may reflect connection interruptions on the participants’ end. The event began at 1:00 PM and ended at 5:00 PM EST.

Fig. 2C shows the number of communications between speakers and the participants. When the event started, there were discussions and questions about the event logistics. In each session, the participants were highly interactive. Communication was minimal around 3:30 PM during a scheduled 30-minute break. When the event restarted after the break, participants’ questions and comments were focused on hands-on training. In addition to the questions and discussion, the participants’ acknowledgments, such as thanking the event organizers, contributed to the spike seen in Figure 2C towards the end of the event. Indeed, the word “Thank” is one of the most frequent words as shown in the word cloud (Fig. 2D). It is no surprise to see words such as “structure”, “protein”, “PDB”, “AlphaFold”, “RoseTTAFold”, “Thank”, and “AI” dominate the word cloud. The word “https” appeared frequently because the panelists, speakers, and the participants often shared links of online resources such as PDB, source codes, GitHub page, *etc*.

### 2.2 Summary of the Presentations and Hands-on Training

The workshop began with an introduction and welcome addresses by Rutgers University faculty and staff members, Stephen K. Burley, IQB Founding Director; J. Barr von Oehsen, Associate Vice President, Office of Advanced Research Computing; and Michael E. Zwick, Senior Vice President, Office for Research. These presenters gave a broad overview on the scope and learning objectives of the crash course and pointed out how this event benefits the research community connected to each of these co-sponsoring organizations.

After the introductory remarks, the experts gave the following presentations: 1) De Novo Protein Structure Prediction: Disruptive Transformation of Biology in 3D by Minkyung Baek (University of Washington), 2) Primer on Artificial Intelligence: Statistical Underpinning and Vocabulary by Sijian Wang (Rutgers University), and 3) Potential for Impact on Research in Cell/Molecular, Cancer, and Structural Biology by Stephen K. Burley (Rutgers University).

Dr. Minkyung Baek, postdoctoral fellow at the University of Washington and main developer of the RoseTTAFold, explained the working principles of AlphaFold2 and RoseTTAFold. Whenever possible Dr. Beak compared and contrasted these two methods. Both methods rely on genomic sequences information and 3D structures from the Protein Data Bank. Initially, the genomics database is used to find the coevolutionary information for amino acid pairs occurring in the sequence to be modeled. The coevolutionary information is useful to find out which amino acid pairs are close to each other in space. From the coevolutionary information, AlphaFold2 and RoseTTAFold can construct contact maps using attention based deep learning methods. In other words, this first step constructs the 2D structure of the protein (contact map) from the 1D information (sequence). Attention based models were successful in Natural Language Processing (NLP) such as BERT and GPT [19]. A second neural network model is trained to predict the 3D structure from the 2D information by matching with the available templates in the PDB. The whole process is repeated until convergence. By design, RoseTTAFold requires fewer iterations compared to AlphaFold2, and, hence, requires less computational resources. With this iterative approach, both methods achieved a remarkable accuracy in predicting 3D structures for proteins. Dr. Baek also highlighted the recent trends in predicting protein multimers. In the initial versions of AlphaFold2 and RoseTTAFold, released earlier in 2021, only monomer predictions were possible. In late 2021, both AlphaFold2 and RoseTTAFold development teams refined their methods to predict structural models for multimeters [20, 21].

Dr. Sijian Wang (Rutgers-New Brunswick Department of Statistics, Associate Professor and IQB Resident Member), presented a primer on AI methods covering the current trends in machine learning and deep learning methods, and highlighted best practices followed in the community. The primer helped the participants understand the difference between supervised and unsupervised machine learning and how they are applicable to classification and regression analysis. In language models, attention based mechanisms have become successful since they are capable of capturing the short-range and long-range connections among the words. Reinforcement learning approaches are becoming popular in modeling intelligent systems in the real world where the agent and environment interact all the time, such as self-driving cars.

Dr. Stephen K. Burley (Director of the RCSB Protein Data Bank, and Rutgers-New Brunswick Department of Chemistry and Chemical Biology University Professor and Henry Rutgers Chair, and IQB Resident Member), highlighted the importance of open data from genomics and protein structures to drive the innovation in biology, and provided example case studies of AlphaFold2 predictions in connection to cell biology, cancer biology, and the structure of SARS-CoV-2 spike protein. Computational models based on AI methods for predicting proteins and protein complexes will likely improve as the PDB grows [7]. In fact, neither AlphaFold2 nor RoseTTAFold would have been possible without the global PDB. Established with just seven protein structures in 1971, today it houses more than 187,000 experimental structures. In future, integrating the models from experiment and computational methods will impact research and education in fundamental biology, biomedicine, bioenergy, and biotechnology/bioengineering.

Following the expert talks, a series of lightning talks highlighted some of the ongoing research projects at Rutgers that applying deep learning (DL) methods. Dr. Guillaume Lamoureux (Rutgers University-Camden, Department of Chemistry Associate Professor) described the application of DL in predicting protein-protein interactions and building shape models and protein docking models. Dr. Vasileios Petrou (Assistant Professor, New Jersey School of Medicine, Department of Microbiology, Biochemistry and Molecular Genetics) explained how AlphaFold2 predictions are becoming an integral part of structure determination using cryo-electron microscopy. Dr. Sagar Khare (Rutgers-New Brunswick, Department of Chemistry and Chemical Biology Associate Professor and IQB Resident Member) described a computational pipeline that used AlphaFold2 to model predicted structures of the Omicron spike protein interacting with the antibodies [16].

The final session was a hands-on tutorial with AlphaFold2 and RoseTTAFold for predicting structures on the campus supercomputer and on-line web servers. The hands-on session covered four possible cases as described in the workflow chart (Fig. 3), and the participants were able to go and do all these four steps.

**Figure 3:**
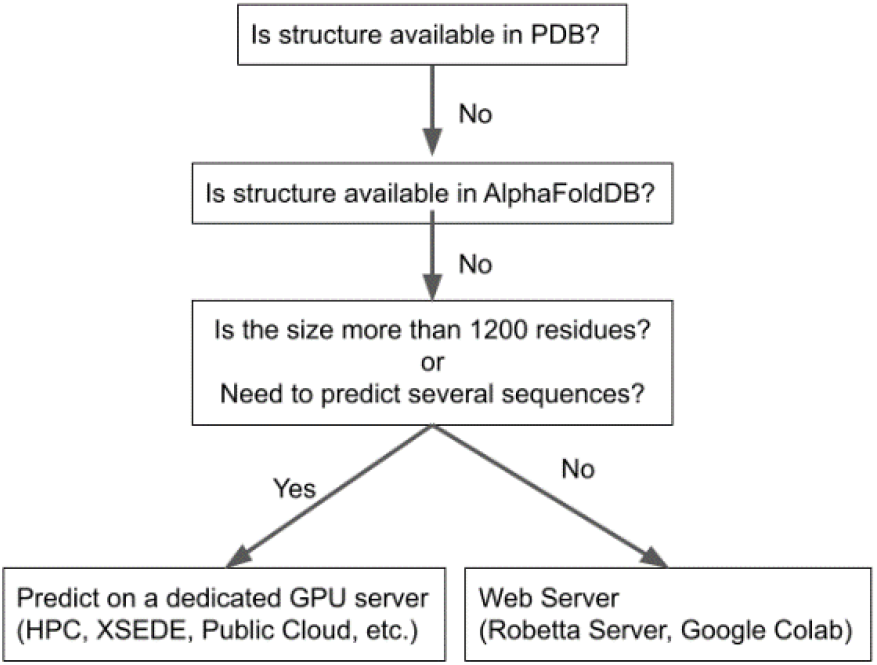
Recommended workflow for obtaining the structure of a protein.

Our workflow (Fig. 3) recommendation was developed for a typical researcher who needs the structure of a protein or protein complex for further investigation or analysis. The fastest and easiest way to obtain the structure of a protein is to look at the existing databases like PDB[18] or AlphaFoldDB[10, 11]. When structure is not available in the existing database, prediction using online servers such as Google Colaboratory (Colab) [22] and Robetta Server [23] is relatively straightforward. This web server approach works well if the size of the protein is less than 1200 residues. Complex tasks, such as modeling of protein complexes or predicting the structure of large proteins, require supercomputing resources that are available in the home institution,, XSEDE [24], or the cloud computing platforms.

We realized that it was challenging to finish all the example calculations within the designated time slot. Although we took a small protein, 1CBN, for our exercise, it still required about 40 minutes to complete on our campus supercomputer. The same calculations would demand more time on the web servers since our jobs would have to wait in the queue. Since the hands-on session was scheduled for only one hour, we required all the participants to submit the jobs on the local campus cluster, Google Colab [22], and Robetta server [23] at the beginning of the hands-on session. While the jobs were running on the servers, we went through the details of the steps outlined in our workflow. This way we optimized the time to conduct the hands-on training. Indeed, a good number of the participants reported that their submitted jobs on the campus cluster finished and produced the desired output files.

## 3 PREPARATIONS FOR THE HANDS-ON TRAINING

In this session, we describe the major steps required in preparing the hands-on training. We built the software stack and the relevant database on the local system and benchmarked the resources requirement for predicting protein structures of various sizes. Based on the benchmark results and the available hardware resources, we finalized the examples for the hands-on training. We developed wrapper scripts to simplify job submissions, executions, and data analysis. We believe that the simplified scripts and examples learned during the hands-on training will serve as templates for the user’s own future calculations and data analysis.

### 3.1. Software and Reference Database

AlphaFold2 offers the option to install the package with a Dockerfile or conda installation script. Both approaches are convenient ways to install packages. Both AlphaFold2 and RoseTTAFold packages come in Python. RoseTTAFold provides the instructions for conda installation. From these instructions, it is straightforward to build a docker or singularity image. In our campus cluster, we made AlphaFold2 and RoseTTAFold in a separate conda environment. We also built an AlphaFold2 stack with the container image.

AlphaFold2 and RoseTTAFold require the reference genomic and protein databases. They are ~ 3 TB in total. The reference data required for RoseTTAFold is the same as that of AlphaFold2. Instead of keeping two copies, we built the database centrally in one location so that AlphaFold2 and RoseTTAFold scripts can utilize the same reference data. Looking from the software release history in the GitHub repositories [26, 27], one can expect that AlphaFold2 and RoseTTAFold will go through rapid updates in terms of algorithm, models, and databases. Building a new version of the software stack with a conda or docker tool does not need a lot of storage or manpower since the installation steps are scripted. However, downloading and managing multiple copies of reference databases require manpower and storage.

### 3.2. Job Execution and Data Analysis

AlphaFold2 and RoseTTAFold provide a Python wrapper script to run the prediction. The script expects multiple input arguments: the name of the fasta file; the location of the database and trained models; the options to control the prediction metrics such as pTM or pLDDT score; and search path for JackHMMER which does the MSA with hidden Markov model. We simplified the job submission by eliminating these arguments for the beginners with the custom wrapper scripts. These scripts and example files are available in the GitHub repository [28]. The other purpose of our wrapper script is to make sure that the jobs are not accidentally executed on the login node. We added a conditional in the script to exit if the job is being executed on the login nodes.

The data analysis of AlphaFold2 and RoseTTAFold methods relies on working with the visualization tools either interactively or through scripts. Our campus cluster supports interactive sessions launched via Open OnDemand [29] [30] or FastX [31]. For structure visualization, we have options such as PyMol, VMD, and Chimera on our local cluster. In addition to structure, the prediction qualities are accessible through the computed metrics such as pLDDT and root mean square deviation (RMSD) values. These metrics can be viewed with any of the plotting tools available in Python, R, or MATLAB. To complete the data analysis, we need to make the participants familiar with these tools which is not practical with the allotted time frame for the hands-on session. Instead of interactively visualizing the live results, we generated the data analysis script that runs on the output files and produces the relevant results as the image files. After the job finished, the participants were able to download the final image files to understand their prediction.

### 3.3. Benchmark Results

The end-to-end calculation of AlphaFold2 and RoseTTAFold demand both memory and compute power. At first, the multiple sequence alignment (MSA) was performed on a given sequence which requires a lot of memory. MSA does not utilize GPUs. After MSA, the deep learning models iteratively refine the structure predictions that demands a lot of compute power. At this stage, GPU accelerators can help speeding up the calculations. Structure predictions with more than one thousand residues require the GPU nodes. For running the crash course, it is not practical to provide the GPUs for all the participants since the participants outnumbered the available GPUs that we can allocate.

Table. 1 shows the benchmark results for running the end-to-end predictions on CPU and GPU compute machines for two proteins. 1CBN is a small protein that has 47 amino acids and 7v7q has 1047 residues for Chain-A. 7v7q, Spike protein of SARS-CoV-2 Delta, is a trimer. For our benchmark, we restricted ourselves to monomer by considering one of the chains. As we pointed out before, MSA does not utilize GPUs. The GPU accelerators impact the compute time required for the model predictions. For example, the GPUs achieved the speedup by a factor of 5 and 20 for 1CBN and 7v7q, respectively. As expected, the newer CPU performed better than the older CPU when we did the tests on 1CBN protein.

**Table 1.** The benchmark results with and without GPU accelerator for 1CBN and 7v7q protein monomers. CPU-A is newer than CPU-B. CPU-A represents Intel Xeon Gold 6130 process, CPU-B represents Intel Xeon E5-2670, and GPU refers to A-100 make.

| PDB ID | # of Amino Acids | MSA (mins) | Model Prediction with CPU-A (mins) | Model Prediction with CPU-B (mins) | Model Predictions with GPU (mins) |
| --- | --- | --- | --- | --- | --- |
| 1CBN | 47 | 30 | 50 | 95 | 10 |
| 7v7q (Chain-A) | 1047 | 140 | 2200 | N/A | 74 |

### 3.4. Web Servers

The RoseTTAFold calculations can be performed on the Robetta server, and the AlphaFold2 calculations can be done in the Google Colab [22]. AlphaFold2 calculations are submitted through an interactive jupyter notebook. These web servers abstract away the complexity of job executions. In the hands-on training, the participants were able to setup the accounts and submit jobs on the Robetta server and Google Colab. While waiting for the calculations to complete, we walked through the results that were completed previously. It is possible to build a workbench solution for AlphaFold2 and RoseTTAFold similar to the work done for the single cell RNA sequence analysis with the Open On Demand approach [32].

## 4 CONCLUSIONS

Our crash course on “Enabling Protein Structure Prediction with Artificial Intelligence” was conducted in collaboration with domain experts and research computing professionals. Led by Rutgers University’s IQB, in collaboration with their OARC, the event was promoted by Rutgers’ Office for Research and the ERN. Since applying AI methods for protein structure prediction is a trending topic, researchers from around the world were eager to learn and apply the AlphaFold2 and RoseTTAFold tools for their research projects. We observed that the participants were enthusiastic about all the presentations and actively engaged via the chat and QAs. Our event attracted more than 750 registrants from around the world.

Although online resources like Google Colab and Robetta servers can enable researchers to carry out protein structure predictions, having AlphaFold2 and RoseTTAFold available on the local campus clusters and national supercomputers like XSEDE will help the community to perform large scale computation and analysis of protein structures. In addition to making the software and relevant reference databases available on the computer clusters, simplified job and data analysis scripts are helpful for the beginners.

Often cutting-edge research topics like AI for protein structure prediction require expertise in multiple fields including structural biology, machine learning, software and data management, and molecular modeling. It is rare for one or two persons to have all the needed experiences in multiple fields. We believe that the approach based on a community wide collaboration involving domain experts and research computing professionals is one of the best ways to build the training programs for the research community in advanced research computing topics.

## ACKNOWLEDGMENTS

The RCSB Protein Data Bank is jointly funded by the National Science Foundation [DBI-1832184, PI: S.K. Burley], the US Department of Energy [DE-SC0019749, PI: S.K. Burley], and the National Cancer Institute, National Institute of Allergy and Infectious Diseases, and National Institute of General Medical Sciences of the National Institutes of Health under grant R01GM133198 [PI: S.K. Burley]. The authors wish to thank all the members of the Eastern Regional Network Structural Biology Working Group for their contributions to the crash course and the ERN Steering Committee for guidance and support. Their work was supported by the National Science Foundation CC* CRIA Eastern Regional Network planning grant (OAC-2018927).

## REFERENCES

1 Jumper, J., Evans, R., Pritzel, A., Green, T., Figurnov, M., Ronneberger, O., Tunyasuvunakool, K., Bates, R., Žídek, A., Potapenko, A., Bridgland, A., Meyer, C., Kohl, S.A.A., Ballard, A.J., Cowie, A., Romera-Paredes, B., Nikolov, S., Jain, R., Adler, J., Back, T., Petersen, S., Reiman, D., Clancy, E., Zielinski, M., Steinegger, M., Pacholska, M., Berghammer, T., Bodenstein, S., Silver, D., Vinyals, O., Senior, A.W., Kavukcuoglu, K., Kohli, P., and Hassabis, D.: ‘Highly accurate protein structure prediction with AlphaFold’, Nature, 2021, 596, (7873), pp. 583–589

2 Baek, M., DiMaio, F., Anishchenko, I., Dauparas, J., Ovchinnikov, S., Lee, G.R., Wang, J., Cong, Q., Kinch, L.N., Schaeffer, R.D., Millán, C., Park, H., Adams, C., Glassman, C.R., DeGiovanni, A., Pereira, J.H., Rodrigues, A.V., Dijk, A.A.v., Ebrecht, A.C., Opperman, D.J., Sagmeister, T., Buhlheller, C., Pavkov-Keller, T., Rathinaswamy, M.K., Dalwadi, U., Yip, C.K., Burke, J.E., Garcia, K.C., Grishin, N.V., Adams, P.D., Read, R.J., and Baker, D.: ‘Accurate prediction of protein structures and interactions using a three-track neural network’, Science, 2021, 373, (6557), pp. 871–876

3 https://www.science.org/content/article/breakthrough-2021#section_breakthrough

4 https://doi.org/10.1038/s41592-021-01380-4, accessed 1 19

5 Hameduh, T., Haddad, Y., Adam, V., and Heger, Z.: ‘Homology modeling in the time of collective and artificial intelligence’, Comput Struct Biotechnol J, 2020, 18, pp. 3494–3506

6 Kuhlman, B., and Bradley, P.: ‘Advances in protein structure prediction and design’, Nature Reviews Molecular Cell Biology, 2019, 20, (11), pp. 681–697

7 Burley, S.K., Arap, W., and Pasqualini, R.: ‘Predicting Proteome-Scale Protein Structure with Artificial Intelligence’, New England Journal of Medicine, 2021, 385, (23), pp. 2191–2194

8 Moult, J., Pedersen, J.T., Judson, R., and Fidelis, K.: ‘A large-scale experiment to assess protein structure prediction methods’, in Editor (Ed.)^(Eds.): ‘Book A large-scale experiment to assess protein structure prediction methods’ (Wiley Online Library, 1995, edn.), pp. ii–iv

9 Jumper, J., Evans, R., Pritzel, A., Green, T., Figurnov, M., Ronneberger, O., Tunyasuvunakool, K., Bates, R., Žídek, A., Potapenko, A., Bridgland, A., Meyer, C., Kohl, S.A.A., Ballard, A.J., Cowie, A., Romera-Paredes, B., Nikolov, S., Jain, R., Adler, J., Back, T., Petersen, S., Reiman, D., Clancy, E., Zielinski, M., Steinegger, M., Pacholska, M., Berghammer, T., Silver, D., Vinyals, O., Senior, A.W., Kavukcuoglu, K., Kohli, P., and Hassabis, D.: ‘Applying and improving AlphaFold at CASP14’, Proteins, 2021, 89, (12), pp. 1711–1721

10 Varadi, M., Anyango, S., Deshpande, M., Nair, S., Natassia, C., Yordanova, G., Yuan, D., Stroe, O., Wood, G., Laydon, A., Žídek, A., Green, T., Tunyasuvunakool, K., Petersen, S., Jumper, J., Clancy, E., Green, R., Vora, A., Lutfi, M., Figurnov, M., Cowie, A., Hobbs, N., Kohli, P., Kleywegt, G., Birney, E., Hassabis, D., and Velankar, S.: ‘AlphaFold Protein Structure Database: massively expanding the structural coverage of protein-sequence space with high-accuracy models’, Nucleic Acids Res, 2022, 50, (D1), pp. D439–d444

11 https://alphafold.ebi.ac.uk/

12 https://oarc.rutgers.edu/

13 https://www.ernrp.org/

14 Jaimes, J.A., André, N.M., Chappie, J.S., Millet, J.K., and Whittaker, G.R.: ‘Phylogenetic Analysis and Structural Modeling of SARS-CoV-2 Spike Protein Reveals an Evolutionary Distinct and Proteolytically Sensitive Activation Loop’, J Mol Biol, 2020, 432, (10), pp. 3309–3325

15 Sternberg, A., and Naujokat, C.: ‘Structural features of coronavirus SARS-CoV-2 spike protein: Targets for vaccination’, Life Sci, 2020, 257, pp. 118056

16 Lubin, J.H., Markosian, C., Balamurugan, D., Pasqualini, R., Arap, W., Burley, S.K., and Khare, S.D.: ‘Structural models of SARS-CoV-2 Omicron variant in complex with ACE2 receptor or antibodies suggest altered binding interfaces’, bioRxiv, 2021

17 https://iqb.rutgers.edu/

18 https://www.rcsb.org/

19 Zheng, X., Zhang, C., and Woodland, P.C.: ‘Adapting GPT, GPT-2 and BERT Language Models for Speech Recognition’, arXiv preprint arXiv:2108.07789, 2021

20 Humphreys, I.R., Pei, J., Baek, M., Krishnakumar, A., Anishchenko, I., Ovchinnikov, S., Zhang, J., Ness, T.J., Banjade, S., Bagde, S.R., Stancheva, V.G., Li, X.-H., Liu, K., Zheng, Z., Barrero, D.J., Roy, U., Kuper, J., Fernández, I.S., Szakal, B., Branzei, D., Rizo, J., Kisker, C., Greene, E.C., Biggins, S., Keeney, S., Miller, E.A., Fromme, J.C., Hendrickson, T.L., Cong, Q., and Baker, D.: ‘Computed structures of core eukaryotic protein complexes’, Science, 2021, 374, (6573), pp. eabm4805

21 Evans, R., O’Neill, M., Pritzel, A., Antropova, N., Senior, A., Green, T., Žídek, A., Bates, R., Blackwell, S., Yim, J., Ronneberger, O., Bodenstein, S., Zielinski, M., Bridgland, A., Potapenko, A., Cowie, A., Tunyasuvunakool, K., Jain, R., Clancy, E., Kohli, P., Jumper, J., and Hassabis, D.: ‘Protein complex prediction with AlphaFold-Multimer’, bioRxiv, 2021, pp. 2021.2010.2004.463034

22 https://colab.research.google.com/github/sokrypton/ColabFold/blob/main/beta/AlphaFold2_advanced.ipynb

23 https://robetta.bakerlab.org/

24 https://www.xsede.org/

25 https://aria2.github.io/

26 https://github.com/deepmind/alphafold

27 https://github.com/RosettaCommons/RoseTTAFold

28 https://github.com/dmbala/AI4Fold_Tutorials

29 Hudak, D., Johnson, D., Chalker, A., Nicklas, J., Franz, E., Dockendorf, T., and McMichael, B.L.: ‘Open OnDemand: A web-based client portal for HPC centers’

30 https://openondemand.org/

31 https://www.starnet.com/fastx/

32 Balamurugan, D., Plazonic, K., Abbey, K., Husain, S., and Syed, N.: ‘Building an Interactive Workbench Environment for Single Cell Genomics Applications’: ‘Practice and Experience in Advanced Research Computing’ (2020), pp. 125–131

